# Architecture and mechanism of a dual-enzyme retron system in prokaryotic immunity

**DOI:** 10.1101/2025.05.23.655875

**Authors:** Xuzichao Li, Qiuqiu He, Yanan Liu, Bin Liu, Zhikun Liu, Hong Chen, Shuqin Zhang, Jie Han, Yingcan Liu, Jie Yang, Hang Yin, Zhenxi Guo, Yong Wei, Zhiyong Yuan, Hongtao Zhu, Heng Zhang

## Abstract

Retrons are bacterial genetic retroelements encoding a reverse transcriptase (RT) and a non-coding RNA (ncRNA)-multi-copy single-stranded DNA (msDNA) hybrid. Diverse effector proteins or domains are found to associate with retrons, typically forming tripartite toxin-antitoxin systems involved in anti-phage defense. Although retrons have attracted growing interest in genome editing technologies, the mechanisms underlying most retron-mediated immune systems remain poorly understood. Here, we characterized a distinct quaternary retron system, Ec78, harboring a dual-enzymatic effector complex, in which the PtuA ATPase and PtuB nuclease act in concert to mediate phage clearance. The cryo-EM structure of the Ec78 complex adopts a flower-basket-like architecture, with two Ec78 retrons engaging the PtuAB effector complexes through a unique msDNA-insertion assembly mechanism. Interestingly, a sensing loop on the RT protein tightly monitors the length of the msDNA, which is likely responsible for phage detection and the subsequent release of the toxic effector complex. We further determined the cryo-EM structure of the retron-unbound effector complex, revealing an arginine-lysine finger loop on the PtuB nuclease that undergoes an ordered-to-disordered transition for enzymatic activation. Together, our work not only delineates the molecular basis underlying the Ec78 system in antiviral defense but also highlights the mechanistic diversity of retron systems in prokaryotic immunity.

## Introduction

Bacterial retrons are specialized genetic elements comprising a reverse transcriptase (RT) and a non-coding RNA (ncRNA)^1^. The ncRNA consists of two parts, msr and msd (referred to msrRNA and msdRNA thereafter). During retron function, the msrRNA forms a scaffold that supports the RT protein, while the msdRNA segment serves as a template for the RT to produce multi-copy single-stranded DNA (msDNA)^2^. Reverse transcription initiates from the 2’-hydroxyl group of a conserved guanosine in the msrRNA, typically resulting in a msrRNA-msDNA hybrid that contains a unique 2’,5’-phosphodiester bond between the branching guanosine and the first nucleotide of the msDNA^3–6^. Since their discovery decades ago, studies of retrons have primarily focused on the mechanism of msDNA biogenesis. Taking advantage of the ability to produce msDNA in situ, retrons have been coupled with CRISPR-Cas9 systems to enable targeted insertion of DNA fragments, offering promising tools for precision genome editing across diverse cell types^7–10^.

However, the biological function of retrons in bacterial remained obscure. In recent years, studies have unveiled that retrons function as components of multigene systems involved in phage resistance, typically consisting of a retron (comprising a reverse transcriptase and ncRNA) and an accessory defense-associated protein or domain, and conferring phage resistance through abortive infection, a common strategy shared by a broad range of defense systems^1,4,11,12^. These systems are widely distributed, classified into nearly 13 types according to the accessory proteins or domains^4^. Despite their widespread distribution and emerging applications, only a few retron systems have been mechanistically characterized. Among them, the Ec86 and Sen2 systems function as tripartite toxin-antitoxin (TA) systems, in which the retron suppress the toxic activity of the associated effectors^13–15^. Structural analysis of the Ec86 interference filament revealed that its msDNA physically cages the N-glycosidase effector in an inactive state. Moreover, effector activation can be triggered by msDNA degradation in the Sen2 system via phage-encoded exonucleases, or msDNA methylation in the Ec86 system, suggesting that modification of msDNA may represent a general strategy for retron activation.

Surprisingly, in contrast to these well-characterized systems which utilize one effector protein for function, the type I-A retron system harbors two enzymatic effector components derived from the Septu system^1,4^. Septu systems are prokaryote defense systems^16^, featuring an ATPase named PtuA and an HNH nuclease named PtuB, and are classified into two types^17^. Type I Septu systems utilize a catalytically inactive PtuA to form a hexamer and recruit two PtuB nucleases for phage defense through degradation of phage DNA^18^. Type II Septu systems, on the other hand, are typically associated with retron genetic elements. Actually, the type II Septu system and the type I-A retron system refer to the same system, but are named differently based on structural or genetic perspectives. However, the mechanism by which the retron cooperates with PtuAB to mediate anti-phage defense remains unclear.

Here, we biochemically and structurally characterized a type I-A retron system, Ec78, and defined it as a quaternary TA system, in which the retron functions as the antitoxin, and the two enzymatic effectors, PtuA and PtuB, form a complex and act in concert to mediate toxicity. We determined high-resolution cryo-EM structures of the two-component effector complex in both retron-bound and free states. Together with biochemical and mutagenesis analysis, we delineated the molecular basis of retron-mediated inhibition, dual-enzymatic effector activation, and their synergistic toxicity, revealing diverse and conserved features among retron-associated defense systems. Our work not only provides mechanistic insights into a dual-enzymatic effector retron system but also offers a foundation for future development of retron-based tools in biotechnological applications.

## Results

### Ec78 system is a toxin-antitoxin system

The Ec78 system belongs to the type I-A retron family and harbors a two-component effector complex derived from the type II Septu system^17^, consisting of an ATPase termed PtuA and an HNH endonuclease termed PtuB (Fig. 1a). Retron genetic elements consist of a RT protein and a ncRNA. The RT protein utilizes the msdRNA as a template for reverse transcription (Fig. 1a). A specialized msDNA molecule is generated and covalently linked to the msrRNA, forming an RNA-DNA hybrid^1^. The ncRNA components in retron systems exhibit considerable variation in both sequence and length, suggesting distinct molecular mechanisms of their nucleic acid elements^4,19^. Therefore, we treated the purified Ec78 complex with proteinase K and RNase A to isolate msDNA for sequencing. DNA sequencing revealed a 78-nucleotide (nt) msDNA synthesized by the RT. We then mapped the sequencing reads along the ncRNA locus and generated msDNA coverage plots for the Ec78 retron (Fig. 1b).

**Figure 1.**
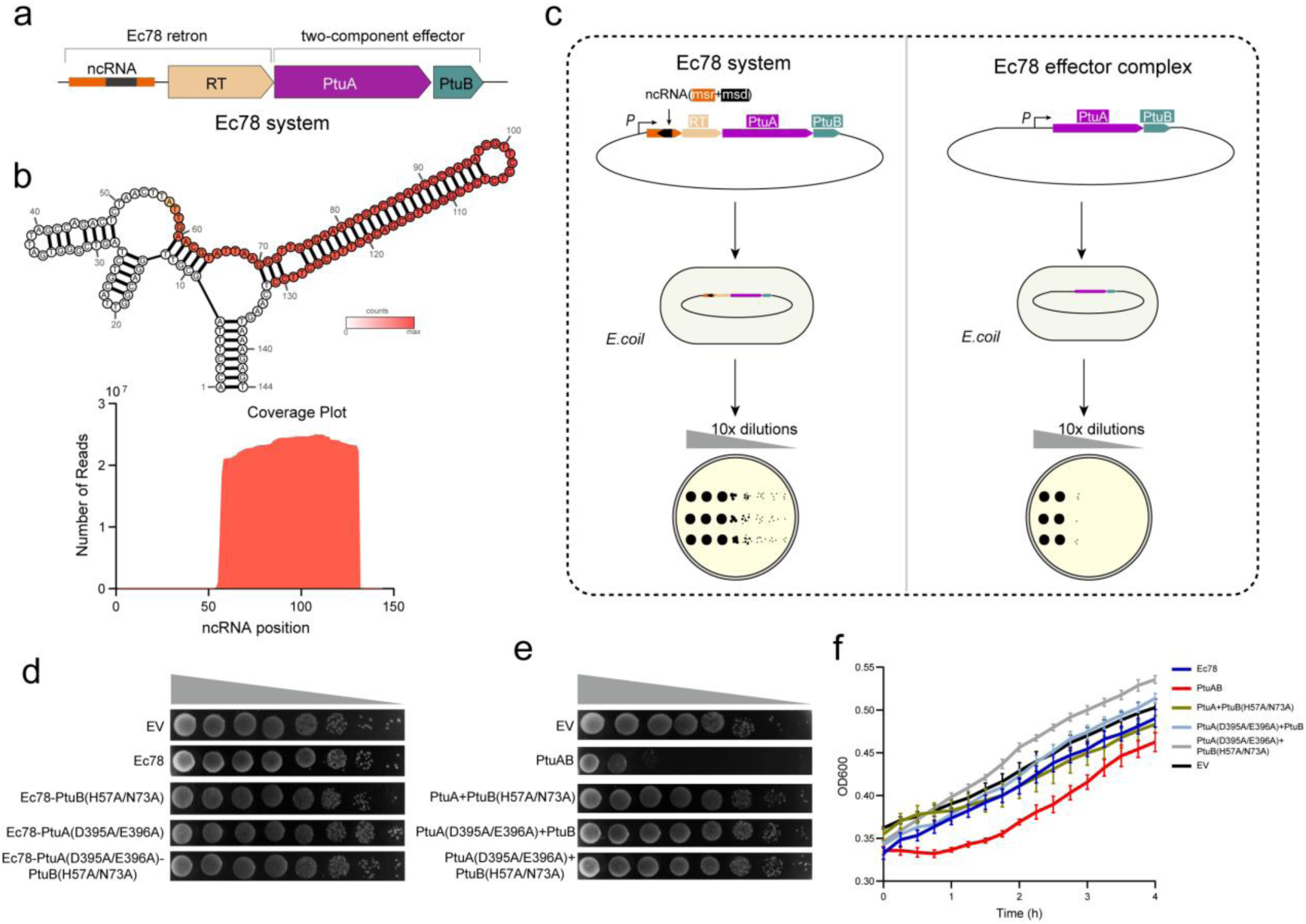
Biochemical characterizations of the Ec78 system. (a) Gene architectures of the Ec78 system from *E. coli* ECONIH5. (b) The msDNA sequencing reads extracted from Ec78 system were mapped to the corresponding ncRNA locus. (c) Schematic diagram of bacterial growth spot assay. Both Ec78 system and Ec78 effector complex were constructed using the native promoter. (d) Bacterial growth spot assay of Ec78 system harboring various mutations in different components. EV, Empty vector. (e) Bacterial growth spot assay of WT or mutant Ec78 effector complex. (f) Growth curves of BL21 (DE3) cells in liquid culture transformed with the WT Ec78 system, empty vector, Ec78 effector complex and effector complex mutants. The results are representative of three individual experiments.

Bacterial retron systems have been reported to function as TA systems, in which both the RT and the msDNA elements act as antitoxins^15^. To investigate whether the Ec78 system is a functional TA system, we cloned the Ec78 system (including retron, PtuA and PtuB) into one plasmid and transferred the vector into *E. coli* cells for expression. We also expressed the Ec78 PtuAB complex or individual Ec78 elements under native promoter (Fig. 1c). Expression of individual components within Ec78 system did not elicit cytotoxicity (Fig. S1). In contrast, expression of the Ec78 PtuAB effectors inhibited *E. coli* cell growth, and co-expression of Ec78 retron rescued the effector-mediated growth arrest, as observed on solid media and in liquid culture (Fig. 1d-f), indicating the Ec78 system as a TA system with PtuAB effector complex function as the toxin. Inducing the catalytically inactivating mutation in the PtuB nuclease completely abolished the toxic activity of the Ec78 effector and restored normal bacterial growth. Alanine substitution of the catalytic residues in PtuA also eliminated the effector-mediated growth arrest (Fig. 1d-f). Hence, both the activity of PtuA and PtuB are essential for the Ec78 effector function.

### Cryo-EM structure of Ec78 system

To elucidate the structural basis of the retron Ec78 system, we co-expressed the Ec78 complex from *E. coli* ECONIH5 and determined its cryo-EM structure at 3.03 Å resolution (Fig. 2a-b, S2). Cryo-EM structures of retron Ec78 system adopts a “flower-basket-like” architecture, comprising four PtuA proteins, one PtuB nuclease, and two Ec78 retrons, with a total molecular weight of 419 kDa (Fig. 2c-f, S2). Two elongated α-helices (termed coiled-coil region) extending from PtuA protomers (A1 and B1) stagger to form the handle of the basket. The remaining portions of the PtuA dimers, together with the RT-msrRNA complex positioned alongside, form the base of the basket, which is split into two units by an extended msDNA stem-loop. Unit A contains a fully assembled Ec78 retron, comprising the RT protein and a hybrid msrRNA-msDNA molecule, forming a “hook-like” architecture. A PtuA dimer of the Ec78 effector faces the Ec78 retron in unit A. In unit B, the msDNA stem-loop of the retron is not involved in complex assembly and remains unresolved in the cryo-EM density, likely due to its intrinsic flexibility. Nevertheless, a complete Ec78 effector complex is clearly resolved, with the PtuB nuclease flanked by PtuA dimer.

**Figure 2.**
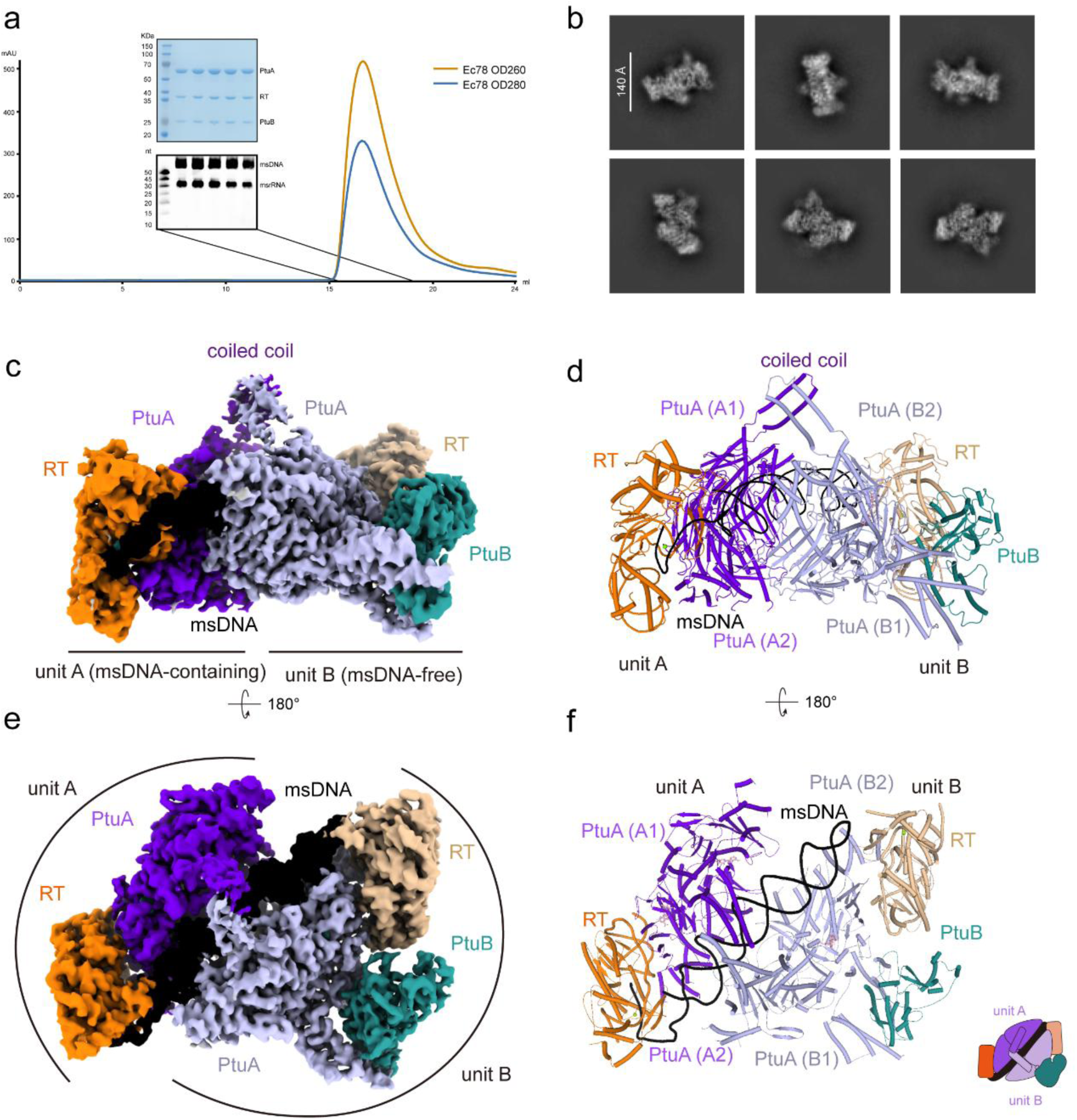
Overall structure of Ec78 system complex. (a) Size exclusion chromatograms of Ec78 system. The peak fractions containing components of Ec78 system were analyzed by SDS-PAGE and urea PAGE. The gels were representative of three repeat experiments. (b) Representative 2D class average with particles of Ec78 complex in different orientations. (c, d) Cryo-EM density map and atomic model of Ec78 system (top view). The ncRNA and msDNA are colored in orange and black, respectively. Protein components are colored in the same scheme as in Figure. 1a. (e, f) Cryo-EM density map and atomic model of Ec78 complex (front view). The same color scheme as Fig. 2c is applied and the corresponding schematic is represented in the lower right corner.

### Structure of Ec78 Retron

High-quality cryo-EM density of unit A enabled unambiguous tracing of most of the Ec78 retron (Fig. 3a-c, S3). In particular, the 311-amino-acid RT protein adopts a canonical right-handed architecture, comprising palm, thumb, and finger domains, and harboring a conserved YADD catalytic core (Fig. 3b-c). A DALI search^20^ of the Ec78 RT protein reveals the highest structural similarity to that in retron Ec86, with RMSD of 3.0 Å, and Z-score of 21.6. Several structural features are unique to Ec78 RT, including two distinctive loop regions (Fig. 3c). Loop I (α6-α7 loop, termed the sensing loop, discussed below), emerges near the finger domain and contacts the distal end of the msDNA stem-loop, and loop II (β5-α8 loop, discussed below) is responsible for interacting with PtuB, suggesting their specialized roles in Ec78 function (Fig. S4a).

**Figure 3.**
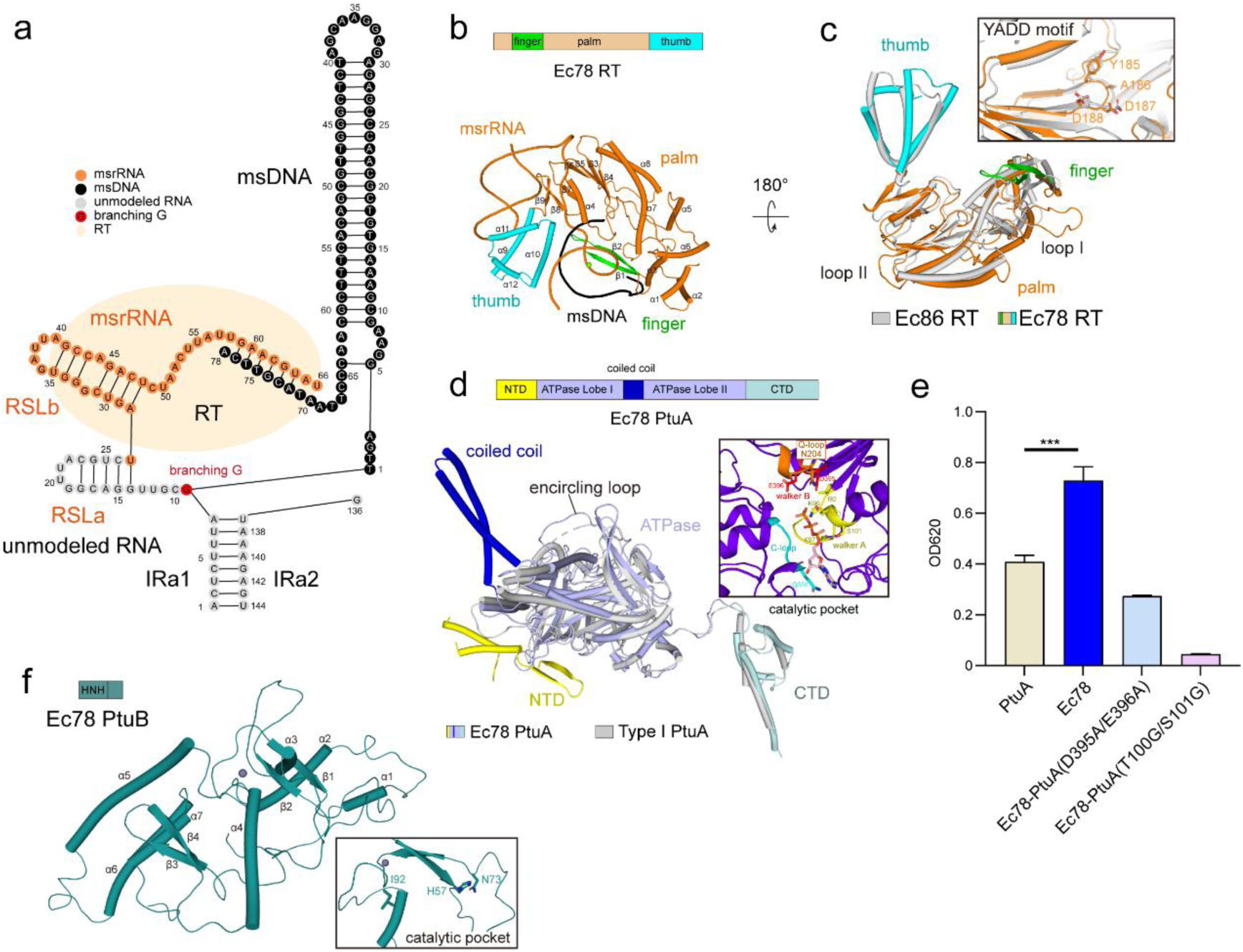
Structural basis of each component in Ec78 system complex. (a) Schematic representation of the msrRNA-msDNA duplex in Ec78 system. Well-resolved nucleotides of msDNA in the duplex (black), msrRNA (orange) and conserved branching G (red) are highlighted, whereas the unresolved nucleotides are indicated by grey circles. (b) Domains distribution of Ec78 RT (upper panel). The atomic model of RT (middle panel) is shown with the same color scheme as the upper panel. Structure alignment of Ec78 RT and Ec86 RT (lower panel). Close-up view of the catalytic YADD motif (inserted panel). (c) Domain organization of Ec78 PtuA (upper panel). The superposition of Ec78 PtuA and Type I PtuA (lower panel). Detailed insight into the catalytic pocket within the ATPase domain is shown in the inserted panel and key residues are shown in stick representation. (d) ATP hydrolysis of PtuA, WT Ec78 system, and Ec78 systems harboring catalytic inactive variants of PtuA (D395A/E396A and T100G/S100G). Three replicates were carried out for each measurement. The data were shown as mean ± SEM. An unpaired Student T-test was used for statistical analysis. Three asterisks indicated P < 0.001. (e) The atomic model of PtuB. Detailed insight into the catalytic pocket is shown in the inserted panel and catalytic residues are shown as sticks.

The msrRNA contains two stem-loop regions, RSLa (14-27 nt) and RSLb (28-48 nt) (Fig. 3a). Only RSLb is resolved in the cryo-EM density, wrapping around the RT thumb domain (Fig. 3b). The inverted repeat sequences (IRa1 and IRa2), located at the 5’-end of the msrRNA and the 3’-end of the msdRNA, along with the branching guanine required for priming msDNA synthesis, remain unresolved in the EM map. Additionally, six nucleotides at the 3’-end of the msDNA, positioned near the RT active site, are base-paired with the msrRNA. The 5’-region of the Ec78 msDNA extends outward to form a nail-like 28-base-pair stem-loop inserting into the entire complex, in stark contrast to the five-pointed star-shaped msDNA structure observed in retron Ec86^21^. Moreover, the two retrons are almost identical to each other except for the msDNA stem loop (Fig. S4b).

### Structure of PtuAB effectors in Ec78 complex

Ec78 PtuA consists of a central ATPase domain, with an additional ear-like domain at its N-terminus and an α-helical bundle domain at its C-terminus (Fig. 3d, S5). The ATPase domain adopts an ellipsoidal bilobed architecture, resembling typical ABC-type ATPases such as Rad50^22^. Lobe I of PtuA ATPase domain is composed of six β-sheets and three α-helices, and folds into an α/β roll structure (Fig. S5a-b). The Lobe II contains five α-helices and six β-sheets and adopts a βα-sandwich fold. The two lobes are connected by a two-helix coiled-coil region (residues 257-305). Additionally, a distinctive encircling loop, spanning approximately 35 residues (residues 204-238), extends from the β8 sheet to the α6 helix in Lobe I and wraps around the surface of the ATPase domain. This structural feature is unique to PtuA and is not observed in other ABC-type ATPases (Fig. 3d, S5c). DALI search indicates that Ec78 PtuA most resembles the PtuA in type I Septu system, with the RMSD of 3.3 Å and DALI score of 30.7. However, the ear-like NTD and coiled-coil region are special in the Ec78 PtuA (Fig. 3d), indicating their potential role in adapting Ec78 function (discussed below). Moreover, the β-hairpin and α-hairpin in the Lobe I and Lobe II of type I PtuA protein involving in its higher oligomerization, are absent in the Ec78 PtuA, suggesting the distinct anti-phage strategies of type I and II Septu systems^18^. The Ec78 complex harbors four copies of PtuA, which are almost identical (Fig. 2c-f, S5d).

Two ATPase domains of Ec78 PtuA dimerize face-to-face, forming two ATP-binding pockets at the dimer interface. Both ATP-binding pockets contain an ATP molecule in unit A, while only one pocket is occupied in unit B and the other site remains open, resulting in a closed conformation of the PtuA dimer in unit A, and a relatively open conformation in unit B (Fig. S5e). Within PtuA dimer, each ATP binding pocket is cooperatively formed by the two PtuA protomers (Fig. 3d). Residues from the Walker A motif (also termed the P-loop) of one PtuA protomer and the signature motif (also known as the C-loop) of the opposing protomer are involved in substrate binding through extensive polar and hydrophobic interactions. Two acidic residues, Asp395 and Glu396, responsible for ATP hydrolysis, are located in the conserved Walker B motif. Moreover, a glutamine residue on the Q-loop typically accommodates the Mg^2+^ ion, which is required for ATPase activity in ABC-type ATPases^22^ (Fig. S5f). In contrast, the corresponding site in the PtuA of the type I Septu system is occupied by a glycine residue (Fig. S5g), resulting in its catalytic inactivity^18^. However, an asparagine residue is present at the corresponding site in Ec78 PtuA (Fig. 3d). Hence, to assess the ATPase activity of PtuA, we performed in vitro ATP hydrolysis assays using PtuA alone and the Ec78 complex. Both the Ec78 PtuA and the Ec78 complex exhibited ATPase activity, confirming that PtuA is catalytically active (Fig. 3e). Mutations in the catalytic residues not only eliminated the ATP hydrolysis but also abolished the effector mediated growth arrest. Taken together, these results further suggest that PtuA is a catalytically active ATPase and its activity is indispensable for effector complex toxicity.

Moreover, one PtuB molecule was observed in the Ec78 complex, comprising an N-terminal HNH nuclease domain and a C-terminal domain. The N-terminal HNH domain adopts a relatively loose conformation, featuring a two-stranded antiparallel β-sheet (β1-β2) flanked by two α-helices (α1-α2) (Fig. 3f, S6), and harbors a catalytic dyad composed of two conserved residues (His57 and Asn73). The C-terminal domain consists of a two-stranded β-hairpin and a four-helix bundle, which are responsible for interaction with PtuA.

### Retron-mediated dual-enzymatic effector complex inhibition

Next, we investigated the assembly of the PtuAB effector complex and how it is neutralized by the Ec78 retron (Fig. 4a-f). In contrast to the previously reported Ec86 system, which contains two complete copies of the effector in each filament segment^13^, the Ec78 complex harbors a single PtuAB effector complex in one of its units, and the other contains only a PtuA dimer without the associated PtuB endonuclease (Fig. 4a). Similar features were also observed in type I Septu system, where three PtuA dimers form a horse-shaped hexamer, but the PtuB is recruited to the two side PtuA dimers^18^. In the Ec78 complex, two α-helical bundle CTDs from the PtuA dimers flank the α5-α7 helices of the PtuB C-terminus, establishing extensive polar and hydrophobic interactions (Fig. 4b). Deletion of the PtuA CTD disrupts PtuB recruitment, impairs effector complex assembly, and consequently abolishes cytotoxicity (Fig. 4f).

**Figure 4.**
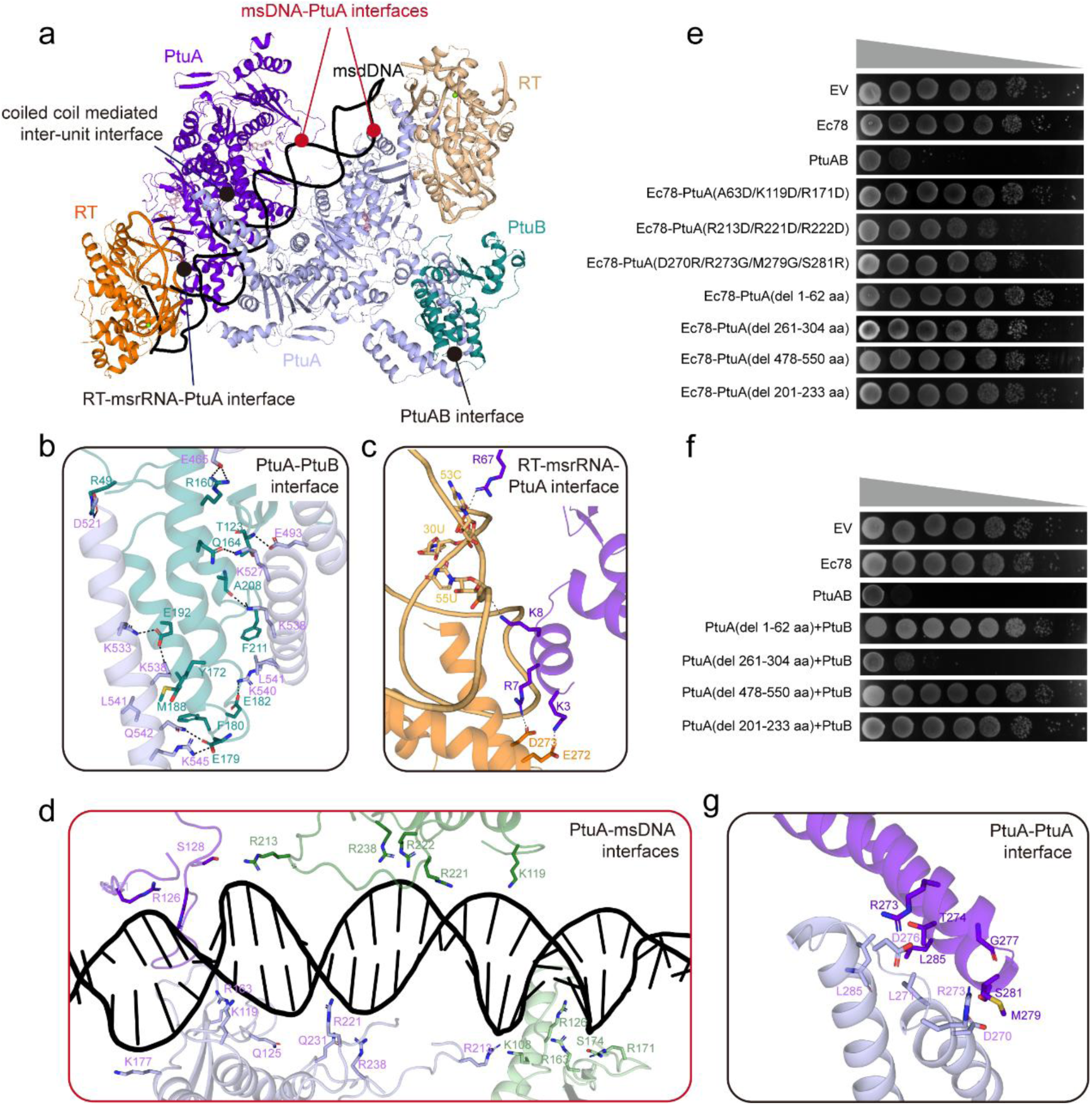
Key interfaces of Ec78 system are essential for functions. (a) Structure of Ec78 complex. Key interaction interfaces are marked using colored circles. (b) Close-up view of the PtuAB interface. Key interaction residues are shown in stick representation. (c) Detailed insight into the RT-msDNA-PtuA interface. Corresponding residues are shown as sticks. (d) The interfaces between the PtuA dimer and msDNA. Key residues that participated in the interaction are shown as sticks. (e) Bacterial growth spot assay of Ec78 systems harboring mutants in Ec78 PtuA. Key structural features on PtuA participate in interacting with retron are deleted to test their importance in effector inhibition. Key msDNA interacting residues are also mutated. (f) Bacterial growth spot assay of Ec78 effector complex harboring truncation mutations within PtuA. Key structural features specific in Ec78 PtuA are mutated, the results indicate their essential role in effector mediated cytotoxity. (g) Close-up view of the ATPase-ATPase interface. Residues involved in interactions are shown in stick representation.

To investigate how the effector complex is anchored by the retron, we analyzed the interactions between the retron and the PtuAB effector complex. Unlike the Ec86 retron system, which utilizes two msDNA molecules to cage the effector for inhibition, the Ec78 system features a long stem-loop structure of the msDNA embraced by two PtuA dimers (Fig. 4a). Additionally, the RT-msrRNA complex caps the side of the ATPase dimer. Together, the retron elements form two major interfaces with PtuA dimers to inhibit the Ec78 effector complex, including the msDNA-PtuA interface and the RT-msrRNA-PtuA interface (Fig. 4c-d).

In particular, the PtuA dimers in units A and B adopt a head-to-head configuration, with the A2 and B2 protomers contacting each other through the α6 helix and β9-β10 loop on the back side (Fig. S7a-b). The front side remains open, forming a positively charged cleft approximately 95 Å in length and 30 Å in width that accommodates the msDNA (Fig. S7c-d). Stretches of positively charged residues on Lobe I of the ATPase domain, especially within the β8-α6 encircling loop, directly engage the stem-loop structure of the msDNA (Fig. 4d). In addition, the ear-like N-terminal domain (NTD) of PtuA, comprising α1-α3 helices and a β1-β2 hairpin, faces the RT thumb domain along with the msrRNA (Fig. 4c, S5a). Positively charged residues within the ear-like region form extensive hydrogen-bonding networks with both the thumb domain and the msrRNA backbone (Fig. 4c).

These structural observations indicate that both the RT-msrRNA and msDNA elements of the Ec78 retron contribute to its antitoxin function. To validate these findings, we generated Ec78 mutants with a deletion in the ear-like NTD, as well as alanine substitutions at msDNA-interacting residues within the PtuA ATPase domain, and expressed these constructs in *E. coli*. As expected, expression of the Ec78 system bearing PtuA mutant R213D/R221D/R222D disrupting the msDNA-PtuA interaction triggered cellular growth arrest, highlighting the essential role of msDNA in effector inhibition (Fig. 4e). Moreover, deletion of the ear-like region and the encircling loop on PtuA resulted in the loss of function in the effector complex, indicating their essential role in effector toxicity (Fig. 4e-f). The retron neutralizes the effector complex by engaging these structural features, thereby inhibiting its toxic activity. Taken together, both the RT-msrRNA and the msDNA elements contribute to effector inhibition by anchoring distinct functional features of PtuA.

More importantly, coiled-coil regions in ABC-type ATPases have been reported to play a crucial role in complex assembly. For instance, the long coiled-coil regions in Rad50/SMC-family ATPases mediate intra-dimer interactions along their distal regions^23^. In contrast, the coiled-coil region in the Aria protein of the PARIS defense system folds into a helical bundle and mediates hexamer assembly through inter-dimer contacts^24^. Distinct from these, the two-helix coiled-coil region in PtuA-A1 protomer bridges with that in PtuA-B1 protomer, mediating inter-unit contacts through both polar and hydrophobic contacts (Fig. 4a, 4g, S7d). The hinge-like arrangement of the coiled-coil regions anchors the msDNA, further stabilizing the PtuA-msDNA assembly (Fig. 4a, S7d). We therefore hypothesized that the assembly of this coiled-coil region in the Ec78 complex may mediate a unique auto-inhibition mechanism of the effector complex. However, deletion of the coiled-coil region in PtuA had no impact on both the effector complex and the Ec78 system (Fig. 4e-f). Moreover, the back sides of the PtuA-A2 and PtuA-B2 protomers are in close contact, which prevents their coiled-coil regions from interacting with each other. As a result, these regions are only visible in the cryo-EM map at a relatively low threshold (Fig. S7e).

Together, the Ec78 retron inhibits the effector complex through both the msDNA and RT-msrRNA elements, which primarily engage the PtuA subunit. Key structural features on PtuA are essential for effector function, and retron closely interacting with these regions, mediating the inhibition of the Ec78 effector complex.

### Activation mechanism of Ec78 effector complex

To further elucidate the molecular basis underlying the activation mechanism of Ec78 effector complex, we attempted to determine the structure of the PtuAB complex. However, we were unable to purify the wild-type PtuAB complex, likely due to its cytotoxicity. We therefore constructed catalytically inactive mutants and purified a PtuAB effector complex bearing T100A/S100A mutation in PtuA. Using cryo-electron microscopy, we determined the structure of the PtuAB effector complex at a resolution of 2.70 Å (Fig. 5a-b, S8). The overall architecture of the effector complex resembles that in the Ec78 system, and the interaction modes between PtuA and PtuB are almost identical (Fig. S9a-b). However, the helix-bundle in the ear-like NTD and the encircling loop in the ATPase domain were not resolved in the PtuAB complex, likely due to increasing flexibility (Fig. 5c). Deletion mutations in these regions totally abolished the toxicity of PtuAB effector complex (Fig. 4f), indicating the essential role of both dynamic regions for effector function, which may be involved in target binding and recognition.

**Figure 5.**
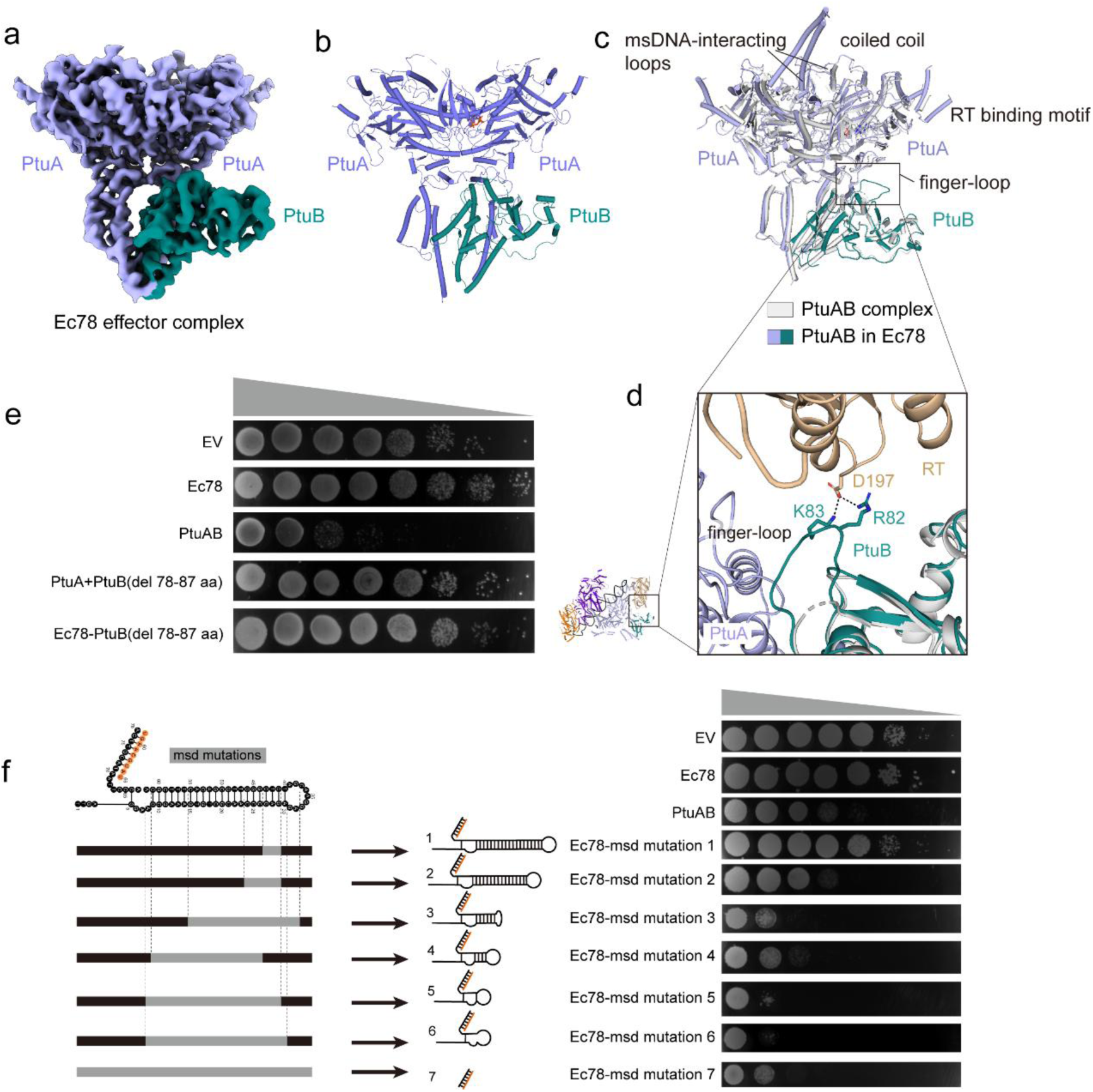
The key finger loop plays an important role in Ec78 system when transitioning from Ec78 complex to Ec78 effector complex. (a) Cryo-EM map of Ec78 effector complex. (b) Atomic model of Ec78 effector complex. (c) Structure comparison between Ec78 effector complex in inhibition and activation states. The arginine-lysine finger loop is marked with a rectangular box. (d) Close-up view of the interaction between the arginine-lysine finger loop of PtuB and RT protein. Key residues are shown as sticks. (e) Bacterial growth spot assay of WT and mutant effector complex. Truncation mutation in the finger loop resulted in the loss-of-function in the effector complex. (f) Bacterial growth spot assay of Ec78 system variants bearing msDNA truncation mutations. The results are representative of three repeats.

Notably, a special arginine-lysine finger loop region (β2-α3, residues 79-90) on the PtuB nuclease exhibited increasing flexibility and could not be resolved in the PtuAB effector complex (Fig. 5c). In contrast, this loop region could be clearly modeled according to high-quality EM density in the Ec78 complex (Fig. 5d). The β5-α8 loop region (loop II) on RT protein contacts and lifts the arginine-lysine finger loop on the PtuB. Specifically, an aspartate residue D197 on the RT loop hydrogen bonds with two basic residues (Arg82 and Lys83) on the PtuB β2-α3 loop, maintaining its extended configuration. Surface electrostatic potential analysis reveals that this arginine-lysine finger loop in PtuB is positively charged, suggesting its potential role in target engagement (Fig. S9c). Deletion of this positively charged loop region abolished PtuAB-mediated growth arrest, confirming its essential role in effector function (Fig. 5e). The observed order-to-disorder transition in this dynamic loop further implies its regulatory role in modulating effector activity. Specifically, the extended conformation of the loop in the Ec78 complex likely obstructs target engagement, thereby suppressing the nuclease activity of PtuB (Fig. 5c-d). Upon dissociation from the retron, the loop adopts a flexible conformation, likely enabling target engagement and subsequently activating nuclease activity. Thus, the arginine-lysine finger loop may function as a molecular “switch” that governs the transition between effector inhibition and activation.

Previous studies have shown that msDNA serves as a sensor of phage infection. Some retron systems, like retron-Sen2, are activated by degradation of msDNA via phage-encoded nucleases^15,25^, suggesting that the reduction in msDNA length may serve as a molecular signal for the activation of retron-associated TA systems. In Ec78, we identified a structural feature that likely responds to this msDNA-reduction signal. Specifically, a loop region on the RT protein (α6–α7 loop, residues 135–155), hereafter referred to as the sensing loop, interacts with the distal end of the msDNA and appears to monitor its length via electrostatic interactions (Fig. S10a). Two basic residues, Lys141 and Arg142, form hydrogen bonds with the phosphate backbone of 36dA and 37dC, respectively. This interaction suggests a ruler-like mechanism, in which the RT sensing loop monitors the msDNA length to regulate downstream effector activation. To test this model, we generated a series of stem-loop truncation mutants, reducing the msDNA stem-loop length to 22, 16, 10, 6, 5, and 0 bp (Fig. 5f). Expression of the Ec78 system with these truncated msDNAs markedly inhibited bacterial growth, particularly in variants with stem-loops shortened to 10 bp or less (Fig. 5f). Conversely, extending the msDNA stem-loop also disrupted proper assembly of the Ec78 complex and resulted in growth arrest (Fig. S10b), reinforcing the importance of precise msDNA length. We further attempted to assess the direct role of the sensing loop by constructing deletion mutations in the sensing loop. However, we failed to construct the plasmids bearing these RT variants, likely due to misfolding or toxicity in *E. coli*, which itself underscores the structural and functional importance of this loop region. Together, these findings support a model in which the msDNA element is essential for Ec78 function, and its precise length is monitored by the RT sensing loop to control effector activation in response to phage infection.

## Discussion

Retrons were discovered more than 40 years ago, but their biological functions remained unclear for decades^2^. Recent studies have revealed that retrons act as anti-phage defense systems, typically through abortive infection (Abi) strategies^1,26^. Retrons are characterized by associated defense components encoded within their gene cassettes and can be grouped into approximately 13 types based on their associated proteins or domains, reflecting considerable mechanistic diversity^4^. However, only a few systems have been well studied^1,15,21^. The type I-A retron systems are uniquely characterized by a dual-enzymatic effector complex^17^. In this study, we resolved high-resolution cryo-EM structures of both the Ec78 system and its isolated PtuAB effector complex. These structures, together with biochemical and mutational analyses, reveal the molecular basis of Ec78 system assembly, effector activation, and phage infection sensing. Our findings support the paradigm that retron systems function as TA systems, in which the retron serves as the antitoxin that neutralizes the toxic activity of associated effectors.

In the Ec78 effector complex, PtuA and PtuB assemble into a 2:1 stoichiometric unit (Fig. 5a-b), in contrast to the 6:2 PtuA:PtuB polymer observed in type I Septu systems^18^. Structural analysis shows that PtuA dimers flank PtuB on both sides, forming a compact and cooperative assembly. Both ATPase and nuclease activities are indispensable for cytotoxicity, indicating that PtuA and PtuB act in concert to execute dual-effector cytotoxicity during abortive infection. Notably, Ec78 PtuA has evolved several unique structural features, including an “ear-like” N-terminal domain (NTD) and an encircling loop, playing critical roles in both effector function and retron-mediated inhibition.

Cryo-EM analysis revealed that the predominant particles of the Ec78 complex adopt a flower basket-like architecture, in which two Ec78 retrons neutralize one complete PtuAB complex and a PtuA dimer (Fig. 2c-f). We also observed particles containing solely retrons and two PtuA dimers (Fig. S2). This observation suggests a dynamic and potentially stepwise assembly process of the Ec78 complex, wherein PtuA dimers are first recruited by the retron and subsequently flank PtuB to complete the effector complex (Fig. 6). Hence, it is conceivable that another PtuB molecule may also be recruited by the other PtuA dimer in unit A, although we were unable to capture Ec78 complex assemblies containing two PtuB molecules, likely due to limitations in the purification strategy.

**Figure 6.**
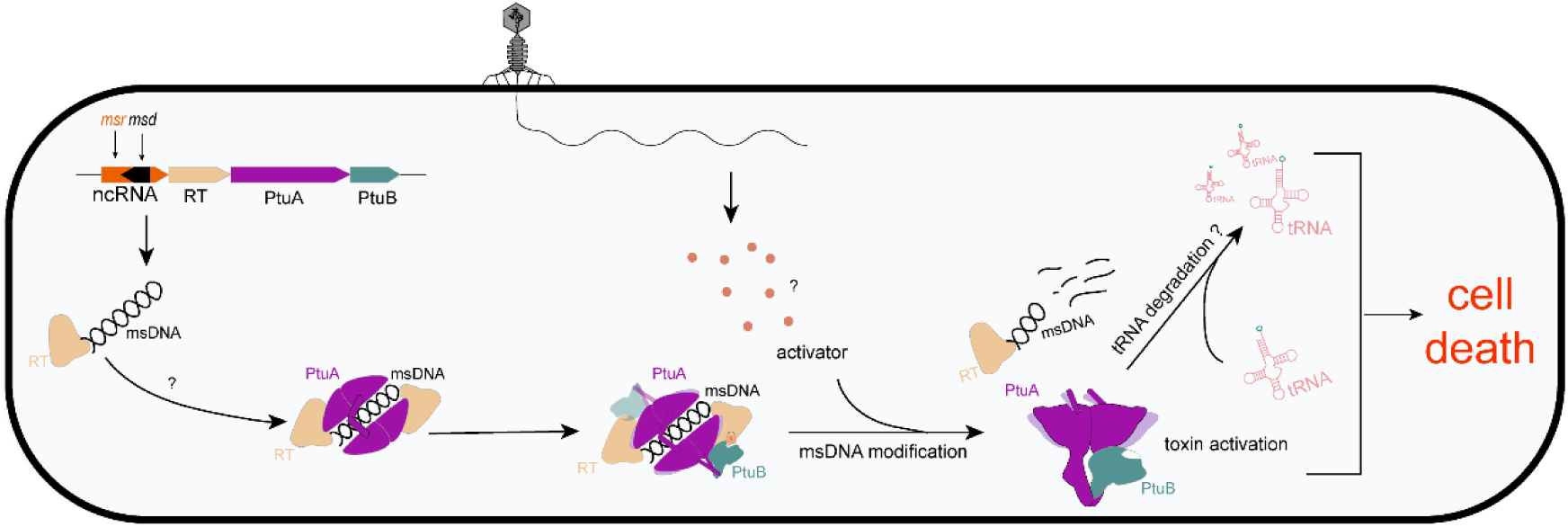
Schematic diagram illustrating the molecular mechanism of Ec78 system.

In line with earlier studies, the Ec78 complex employs both RT and msDNA components to inhibit effector activity. However, the detailed structural basis for toxicity inhibition is notably distinct. In the Ec78 complex, a nail-shaped stem-loop of msDNA is embraced by two PtuA dimers (Fig. 4a), in contrast to the Ec86 filament, where two msDNA molecules cage the effector dimer inside. The RT and msDNA elements tightly contact the PtuA dimers, capping their unique structural features essential for effector toxicity, thereby neutralizing the effector complex in an inhibited state (Fig. 4a-d). Moreover, the catalytic dyad of the PtuB nuclease is solvent-exposed in the Ec78 complex, probably indicating an active state. However, a dynamic arginine-lysine loop, essential for PtuB activity, is fixed by the RT palm domain, thereby suppressing the PtuB nuclease activity (Fig. 5d).

msDNA modification has been reported as a common strategy for toxin activation. Building on previous evidence that the phage-encoded nuclease may serve as an activator of the Ec78 system^25^, we found that introducing msDNA truncation mutations into the Ec78 complex resulted in growth arrest in *E. coli*, highlighting the importance of msDNA in Ec78 function and supporting a potential activation mechanism through msDNA degradation (Fig. 5f). More interestingly, we identified a functional unique loop in the RT palm domain that senses msDNA length (Fig. S10a). This sensing loop recognizes the distal end of the msDNA stem-loop and functions as a molecular checkpoint that monitors msDNA integrity, likely stimulating the release of the effector and facilitating toxicity activation upon phage infection. Upon release of the effector complex from the Ec78 retron, the positively charged structural features on PtuA and the arginine-lysine finger loop on PtuB undergo order-to-disorder transitions, which likely enables target engagement (probably tRNA as previously reported^25^), thereby activating the toxin (Fig. 6). Altogether, our study not only provides in-depth insights into retron-mediated defense mechanisms but also paves the way for the future development of retron-based tools in biotechnology applications.

## Methods

### Plasmid construction

The genomic locus of the Ec78 system (GenBank: CP026202.1) including the ncRNA, RT, PtuA, and PtuB coding genes, was synthesized (Sangon) and cloned into appropriate vectors. The co-expression plasmids for Ec78 system, excluding the ncRNA (RT, PtuA and PtuB) were cloned into 2HR-T vector (Addgene #29718) using the native genomic locus sequence, with a His-strep tag attached to the N-terminus of RT, all under the control of a single T7 promoter. The ncRNA coding gene was cloned separately into the 13S-A vector (Addgene #48323). These two plasmids were co-transformed to express the full Ec78 system complex.

The Ec78 system plasmid and the plasmid containing Ec78 effector complex were both inserted into pACYCDuet-1 (Addgene #128837) and pBAD LIC vectors (Addgene #37501), and were independently transformed into *E. coli* BL21 (DE3) cells for further experiments. The point mutations were generated by QuickChange mutagenesis (Takara) following the manufacturer’s instructions.

### The msDNA extraction and sequencing

The purified Ec78 complex was treated with proteinase K and the inactive protein complex was then incubated with RNase A to digest RNA. The purified ssDNA was processed using the Vazyme ssDNA Library Prep Kit (ND620) following the protocol to prepare the sequencing library. The ssDNA sequencing was conducted at GENEWIZ.

### Protein expression and purification

The BL21(DE3) competent cells were transformed with expression plasmids and plated on LB agar containing appropriate antibiotics. Single colonies were picked and inoculated into 5 mL of starter culture, grown overnight at 37°C, and subsequently transferred to a 1 L culture. Protein expression was induced with 0.2 mM isopropyl-β-D-thiogalactoside (IPTG) when the optical density at 600 nm reached ∼0.6. Cultures were grown at 16°C with orbital shaking at 160 rpm for 16 h.

Cells were harvested by centrifugation at 4°C for 10 min, and the resulting pellets were resuspended in binding buffer (25 mM HEPES pH 8.0, 500 mM NaCl, 2 mM β-mercaptoethanol). The cell suspension was lyzed by sonication, and the lysate was centrifuged to remove debris. The supernatant was incubated with Ni-NTA, Strep-Tactin, depending on the tag of the target protein. Proteins were eluted with binding buffer supplemented with 300 mM imidazole, 5 mM biotin, respectively.

The eluate was further purified by size-exclusion chromatography, with or without prior ion-exchange chromatography. Fractions were analyzed by SDS-PAGE to confirm protein components and by Urea-PAGE to detect nucleic acid components when present. Peak fractions with the correct size of proteins and/or nucleic acid complexes were concentrated, aliquoted and stored in −80°C prior to use.

### ATPase assay

The ATPase assay was performed in a reaction buffer containing 100 mM Tris-HCl (pH 7.5), 600 mM NaCl, 20 mM MgCl_2_ and 8 mM DTT. Purified target proteins were incubated with ATP at a final concentration ratio of 1:40 in the reaction buffer. The total volume of the reaction was 80 μl. The mixture was then incubated at 37 ℃ for 60 min. ATP hydrolysis was determined by measuring the absorbance at 620 nm using a microplate reader (BioTek Synergy H1 hybrid). The data were presented as mean ± standard error of the mean (SEM). An unpaired Student T-test was used for statistical analysis. A significance threshold of P < 0.001 was applied, denoted by three asterisks in Figure 3e.

### Bacterial growth spot assay

Both Ec78 system plasmid and the plasmid containing Ec78 PtuAB complex were constructed using a native promoter, and the two recombinant plasmids were transformed into *E. coli* for amplification. Cells were cultured until the optical density at 600 nm reached ∼0.3. After equal proportion dilution, these dilutions were spotted onto LB agar plates containing the appropriate antibiotic. The plates were incubated overnight at 37°C and were imaged using a gel imager (Azure Biosystems).

For msDNA mutations, the plasmids used the pBAD promoter and were transformed into *E. coli.* Individual colonies were grown in the liquid culture containing 0.4% glucose and antibiotic, and spotted in LB agar plates in the presence of 0.2% arabinose and appropriate antibiotic.

### Bacterial growth assay

*E. coli* cells with the Ec78 system, mutations, and Ec78 PtuAB complex were cultured at 37°C. Then, 180 μl of culture was mixed with 20 μl of cells were transferred into a 96-well plate with the appropriate antibiotic and incubated at 37°C with shaking. The OD600 of the cells was measured using a microplate reader (Biotek) every 15 min for a total duration of 4 h.

### Cryo-EM sample preparation and imaging

For both Ec78 system complex sample and Ec78 effector complex sample, an aliquot of 3.5 μl purified target protein complex at 1∼2 mg/ml was applied to glow-discharged Au R1.2/1.3 holey carbon girds (300 mesh, Quantifoil). After a 5 s incubation, grids were blotted for 2 s at a blot force setting of 2. Then the grid was plunge-frozen into liquid ethane, which was pre-cooled by liquid nitrogen, using Vitrobot Mark IV (FEI Thermo Fisher) at 4°C and under 100% humidity.

Both the datasets of the Ec78 system complex and the Ec78 effector complex were collected on a FEI Titan Krios, equipped with a Falcon4 detector at an acceleration voltage of 300 kV, and at a nominal magnification of 165,000× with a calibrated physical pixel size of 0.74 Å/pixel. Cryo-EM images were collected by SerialEM^27^ or EPU^28^ with the defocus range of −1.2 to −1.6 μm. Each micrograph was dose-fractioned into 32 frames. The total dose was 50.0 e^-^/Å^2^ per micrograph.

### Cryo-EM data processing

For the dataset of Ec78 system complex, 5,499 micrographs were collected and subsequently processed in CryoSPARC^29^, with the beam-induced motion by MotionCor2^30^ and CTF estimation^31^. Micrographs with an estimated CTF resolution higher than 5 Å were excluded, resulting in 4,892 micrographs retained for further analysis. Particles were picked independently using blob picking, template picking and Topaz picking^32^, respectively. After removing duplicate particles from the three particle stacks, a final set of 1,557,724 particles was generated. Then multiple rounds of 2D classification were carried out, yielding 400,145 promising particles for ab-initio reconstruction. After obtaining volumes, multiple rounds of heterogeneous refinement and 3D classification were employed to refine these volumes. Particles re-extraction, local CTF refinement and non-uniform refinement were performed to improve the volume quality, and the final resolution of the best volume was at 3.03 Å.

For the dataset of Ec78 effector complex, 7987 micrographs were collected and processed in CryoSPARC. Motion correction and CTF estimation were performed, followed by manually curate exposures. 7937 micrographs were used for particle picking utilizing blob picking and template picking. Particles with protein features were selected to train a Topaz model. After Topaz extraction, 1,681,139 particles were picked. Multiple rounds of 2D classification were then performed to eliminate junk particles. Ab-initio reconstruction generated the initio volumes and heterogeneous refinement was further conducted to screen good particles. Next, another round of Ab-initio reconstruction and non-uniform refinement were carried out, generating the final map at 2.70 Å.

### Model building and refinement

For the Ec78 system complex, the initial atomic models of retrons, PtuA and PtuB subunits predicted by Alphafold3^33^ were fitted manually into the cryo-EM density map using ChimeraX^34^. For modelling of the nucleotides, docked subunits were used to segment the density map and the nucleotides were built into the remaining density manually in Coot^35^. Then the nucleotides were manually refined to trace the density and map to the sequence. Further refinements were performed by *phenix.real_space_refine*^36^. For the Ec78 effector complex, the protein subunits from the refined structure of Ec78 system complex were separately docked into the EM density in ChimeraX. Further iterative refinements were performed by *phenix.real_space_refine*. The quality of all the final models was validated by MolProbity in Phenix^37^. Data collection and model refinement statistics are concluded in Supplementary Table 1.

## Data availability

The atomic coordinates have been deposited in the Protein Data Bank under accession codes 9L7P (Ec78 system complex) and 9U9Y (Ec78 effector complex). Cryo-EM maps have been deposited in the Electron Microscopy Data Bank under corresponding accession codes EMD-62876 and EMD-63972.

## Acknowledgements

We thank the Core Facility of Research Center of Basic Medical Sciences in Tianjin Medical University for providing technical assistance. We gratefully acknowledge ZQ.Guo and HS.Li (Shuimu Biosciences), DP. Sun (Institute of Physics, Chinese Academy of Sciences), CD.Qin (Cryo-EM Platform, School of Life Sciences, Peking University), and X. Wang and C. Zhang (Cryo-EM Facility, Changping Laboratory) for their expert assistance in cryo-EM data collection. This work was supported by the National Natural Science Foundation of China (32322040 to H.Zhang and 32471015 to H.Zhu), Chinese Academy of Sciences (E2VK311RA1 to H. Zhu), the Natural Science Foundation of Tianjin Municipal Science and Technology Commission (23JCZDJC00410), The Foundation of Tianjin Science and Technology Commission (23ZYCGSY00750) and Scientific Research Program of Tianjin Municipal Education Commission (2023ZD011 to H.Z. and 2024ZD037 to J.Y., and 2022KJ192 to H.Y.)

## Author contributions

Conceptualization: Z.Y., H.Zhu, and H.Zhang; Experimental studies: Q.H., Yanan L., B.L., Z.L., H.C., S.Z., J.H., and J.Y.; Data analysis: X.L., Q.H., Yanan L., Z.L., S.Z., J.H., Yingcan L., J.Y., H.Y., Z.G., Y.W., H.Zhu and H.Zhang; Supervision: H.Zhang; Manuscript writing: X.L., Z.L. and H.Zhang with contributions from all authors.

